# RERconverge: an R package for associating evolutionary rates with convergent traits

**DOI:** 10.1101/451138

**Authors:** Amanda Kowalczyk, Wynn K Meyer, Raghavendran Partha, Weiguang Mao, Nathan L Clark, Maria Chikina

## Abstract

**Motivation:** When different lineages of organisms independently adapt to similar environments, selection often acts repeatedly upon the same genes, leading to signatures of convergent evolutionary rate shifts at these genes. With the increasing availability of genome sequences for organisms displaying a variety of convergent traits, the ability to identify genes with such convergent rate signatures would enable new insights into the molecular basis of these traits.

**Results:** Here we present the R package RERconverge, which tests for association between relative evolutionary rates of genes and the evolution of traits across a phylogeny. RERconverge can perform associations with binary and continuous traits, and it contains tools for visualization and enrichment analyses of association results.

**Availability:** RERconverge source code, documentation, and a detailed usage walk-through are freely available at https://github.com/nclark-lab/RERconverge. Datasets for mammals, *Drosophila*, and yeast are available at https://bit.ly/2J2QBnj.

**Supplementary information:** Supplementary information, containing detailed vignettes for usage of RERconverge, are available at Bioinformatics online.

## 1 Introduction

A major motivation in evolutionary biology is to determine which genetic changes underlie phenotypic adaptations. Convergent evolution, in which the same phenotype arises independently in distinct evolutionary lineages, provides natural replicates of phenotypic adaptation that aid researchers in linking phenotypes to their underlying genetic changes. Selection repeatedly targets the same genes in several known cases of phenotypic convergence, including Prestin in echolocation in bats and marine mammals (Li *et al.*, 2010), *Mc1r* in reduced pigmentation in multiple vertebrate species (Kronforst *et al.*, 2012), and *Na_v_* 1.4 in toxin resistance in snakes (Feldman *et al.*, 2012). This pattern leads to the prediction that genes responding selectively to a convergent environmental or phenotypic transition may show convergent shifts in evolutionary rates (i.e., number of nucleotide or amino acid substitutions per unit time), due to reduced or increased selective constraints on those genes. For example, both genes and non-coding elements involved in eye function lose selective constraint in subterranean mammals, and the corresponding rate shift signal can be used genome-wide to identify candidate regions with eye-specific function (Partha *et al.*, 2017). Thus, evolutionary rates provide a rich source of genome-wide molecular data that, in combination with convergent phenotypes, can be used to infer targets of convergent selection.

Several research groups have developed methods to identify patterns of convergent evolutionary rate shifts associated with convergent changes in traits or environment, including Coevol (Lartillot and Poujol, 2011), Forward Genomics (Hiller *et al.*, 2012; Prudent *et al.*, 2016), and PhyloAcc (Hu *et al.*, 2018). However, existing methods to identify these patterns still have several areas for improvement, including support for both binary and continuous traits (Kowalczyk *et al.*, prep), compute times that enable genome-wide analyses, robust statistical treatment (Partha *et al.*, prep), and ease of visualization and downstream analyses. To address these issues, here we present RERconverge, a software package that implements tests of association between evolutionary rates of genetic loci and convergent phenotypic traits, both binary and continuous (pipeline illustrated in Figure 1). RERconverge incorporates and corrects for known phylogenetic relationships among included species, performs rapid analyses of genome-scale data, and implements appropriate statistical techniques to reduce the effect of outliers and correct for multiple testing. This easy-to-implement software enables biologists with extensive organismal knowledge to apply computational techniques to quickly test hypotheses and link molecular data with phenotypes.

**Fig 1.**
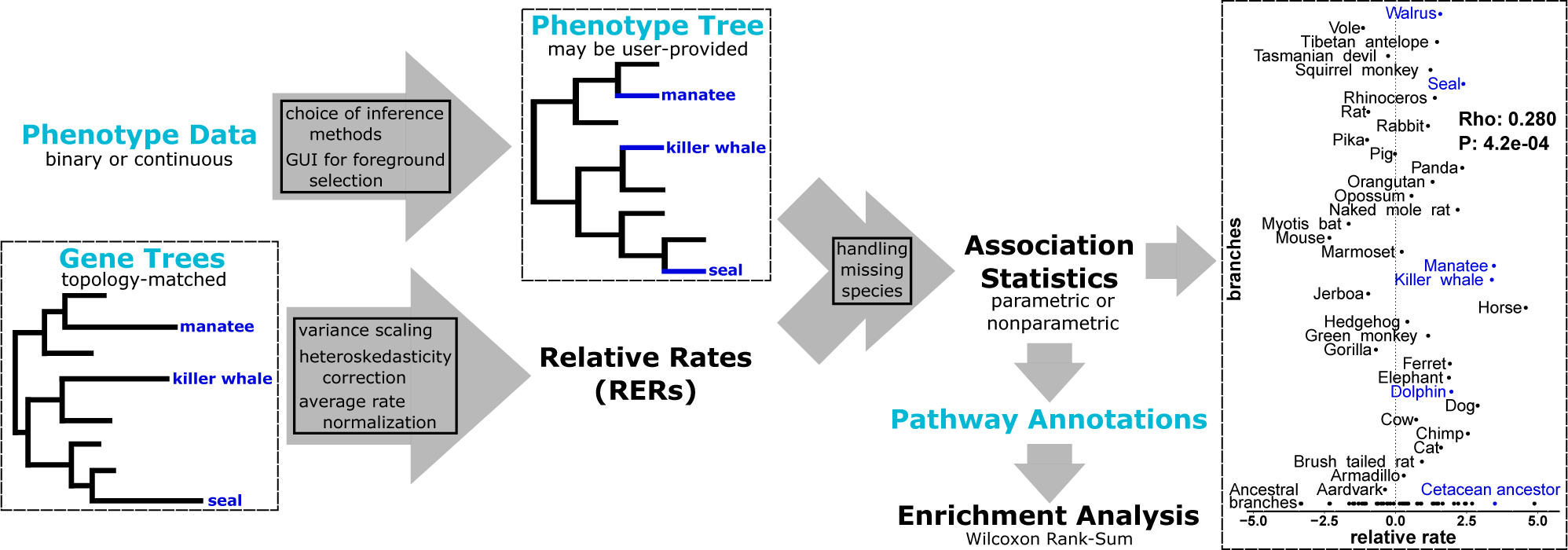
Schematic of the RERconverge pipeline focusing on the example discussed in the case study. Far right figure shows RER for olfactory receptor *OR9Q2*, a top hit for marine-specific acceleration. For additional details of the methods (boxes in arrows), see (Partha *et al.*, prep).

## 2 Description and implementation

### 2.1 Basic usage of RERconverge

The RERconverge package runs within an installation of the R software (R Core Team, 2017) on any platform (Linux, Windows, Mac OS). The user provides the following data as input:

- A set of phylogenetic trees, all with the same topology, with branch lengths for each tree calculated from alignments of that “gene” sequence (these can also represent non-coding sequences such as enhancers) using software such as PAML (Yang, 2007).
- Values for a trait of interest for extant species/tips of the tree. RERconverge can infer values for internal branches of the tree, or the user can provide these values via a single tree or by using RERconverge interactive branch selection to select foreground branches.

Detailed instructions for RERconverge are available within the Supplementary information to this article or at https://github.com/nclark-lab/RERconverge/wiki/Vignettes.

### 2.2 Rapid estimation and visualization of Relative Evolutionary Rates (RER)

RERconverge offers efficient computation of gene-specific rates of evolution on branches of phylogenetic trees in genome-scale datasets. These gene-specific rates of evolution, termed relative evolutionary rates (RER), reflect the amount of sequence divergence on a particular branch after correcting for non-specific factors affecting divergence on the branch such as time since speciation and mutation rate. Additionally, using a combination of statistical approaches including data transformation and weighted linear regression, RERconverge provides estimates of relative evolutionary rates that are robust to several factors introducing outliers in the dataset, such as the presence of distantly related species in the phylogeny (Partha *et al.*, prep). To accelerate the computations underlying RER calculations, key functions are written in C++ and integrated with the R code through the Rcpp package (Eddelbuettel and François, 2011).

### 2.3 Flexible specification of binary and continuous trait evolution

For binary traits, RERconverge provides multiple methods for users to specify which branches are in the “foreground”, meaning those that display the convergent trait of interest. Users can directly supply foreground branch assignments by providing a phylogenetic tree with appropriate branch lengths or by interactively selecting branches on a phylogeny. Alternatively, users can specify which extant species are foreground and allow the software to infer which internal branches to consider foreground under a variety of scenarios. These include whether to consider only branches at a transition to a convergent phenotype as foreground or to also consider subsequent branches that preserve the phenotype as foreground, whether the foreground trait can be lost or only gained, and whether each foreground branch should be weighted equally.

In addition to supporting analysis of binary traits, RERconverge can perform correlation analysis between evolutionary rates and continuous traits (Kowalczyk *et al.*, prep). Trait evolution is modeled on a trait tree constructed from user-provided phenotype values for extant species provided by the user and ancestral nodes inferred through a phylogenetic model. By default, trait tree branch lengths represent change in the trait between a species and its ancestor, which removes phylogenetic dependence between branches. For completeness, other options allow branch lengths to be calculated as the average or terminal trait value along a branch to represent the state of a trait rather than its change.

### 2.4 Genome-wide association between RER and traits, with correction for multiple testing

RERconverge rapidly computes the association between gene-specific relative evolutionary rates and the user-specified traits for large sets of genes. The program estimates the correlation between gene tree branch lengths and the values for traits along those branches using (by default) Kendall’s Tau for binary traits and a Pearson linear correlation for continuous traits. Full gene lists are returned with correlation statistic values, the number of data points used for the correlations, and adjusted p-values (FDR) derived using the Benjamini-Hochberg correction (Benjamini and Hochberg, 1995).

### 2.5 Assessing gene set enrichment in results

Further analysis of gene lists from correlation analyses can be performed using built-in RERconverge pathway enrichment functions. These functions perform a rank-based enrichment analysis by using a Wilcoxon Rank-Sum Test to detect distribution shifts between a subset of genes and all genes in annotated pathways. Enrichment results are returned with pathway names, enrichment statistics, Benjamini-Hochberg corrected p-values, and ranked pathway genes.

## 3 Case study

We demonstrate the features of RERconverge using gene trees derived from the coding sequences of 19,149 genes for 62 mammal species, along with foreground branches that represent lineages of the mammalian phylogeny whose members live predominantly in marine aquatic environments (Chikina *et al.*, 2016; Meyer *et al.*, 2018; Partha *et al.*, prep). For this dataset, RERconverge ran in 1 hour and 18 minutes on RStudio with R version 3.3.0 installed on a computer running Windows 10 with 16 GB of RAM and an Intel Core i7-6500U 2.50GHz CPU. Top categories of functional enrichment included olfactory transduction (FDR=7.2e-90), GPCR signaling (FDR=1.4e-38), and immune functions (FDR=5.4e-7), as in Chikina *et al.* (2016). We show the RERconverge workflow and visualization of RERs for a top olfactory receptor, *OR9Q2*, in Figure 1.

## 4 Summary

RERconverge is an easy-to-implement method that can quickly test for associations between genes’ relative evolutionary rates and traits of interest on a phylogeny. This enables researchers studying a wide variety of questions to generate lists of candidate genes associated with evolutionarily important traits and to explore through enrichment analyses the biological functions showing the most evidence of molecular change in association with these traits.

## 5 Acknowledgments

We thank Andreas Pfenning, Qingyuan Zhao, Ben Rubin, and Tim Sackton for feedback on earlier versions of this software, as well as all members of the Chikina, Clark and Kostka labs for helpful discussion. This project was supported by NIH R01 HG009299-01A1 to N.L.C. and M.C. A.K. was supported by NIH T32 training grant T32 EB009403 as part of the HHMINIBIB Interfaces Initiative.

